# LIMD1 Loss as an Early Driver of PD-L1 Upregulation and Immune Evasion in Lung Cancer

**DOI:** 10.1101/2025.02.06.636805

**Authors:** Kunal M. Shah, Paul T. Kennedy, James R. M. Black, Kevin Litchfield, Krupa Thakkar, Maria F. Contreras-Gerenas, Kirsten Brooksbank, Oliver Yuan, Paul Grevitt, Sarah Charrot, Jeff Davies, Lekh N. Dahal, Dimitris Lagos, Nicholas McGranahan, Tyson V. Sharp

## Abstract

*LIMD1* is a tumour suppressor gene frequently lost in non-small cell lung cancer (NSCLC), but its role in cancer-immune cell interactions remains unexplored. Here, we demonstrate that LIMD1 loss results in upregulation of the key immune checkpoint protein PD-L1. Using multi-region sequencing from the TRACERx dataset, we identify that LIMD1 loss is clonal in over 80% of squamous cell carcinoma (LUSC) and 40% of lung adenocarcinoma (LUAD) cases, correlating with increased PD-L1 expression. LIMD1 deficiency results in upregulation of basal and IFNγ-induced PD-L1 expression in NSCLC cells and, consistent with its early loss during oncogenesis, in primary human small airway epithelial cells. Mechanistically, we demonstrate that LIMD1 interacts with the E3 ubiquitin ligase ARIH1 to mediate efficient PD-L1 ubiquitination and degradation, a process that is significantly impaired in LIMD1-deficient cells, resulting in increased PD-L1 stability. As a consequence, LIMD1-deficient tumour cells suppressed CD8+ T cell activation *in vitro*, and blockade of PD-L1 reversed this suppression. Clinically, we show that LIMD1 loss is associated with enhanced response to immune checkpoint inhibitors (ICIs) in NSCLC patient cohorts, revealing a novel cancer cell-intrinsic correlation of ICI efficacy. Our results uncover a tumour suppressor-mediated mechanism of PD-L1 expression and pave the way for stratified immunotherapy approaches in *LIMD1^-/-^* NSCLC.

## Introduction

Lung cancer is responsible for 21% of total cancer deaths, with approximately 35,000 deaths annually in the UK ^1^. This high mortality rate is attributed to late diagnosis and frequent therapeutic resistance. Although immune checkpoint therapies have transformed cancer treatment, response rates remain variable, typically around 20-40% ^2^. Enhancing these response rates necessitates a comprehensive understanding of the molecular mechanisms governing immune checkpoint protein expression.

The programmed death-1/programmed death ligand-1 (PD-1/PD-L1(CD274)) axis is an immunosuppressive pathway that regulates cytotoxic T cell responses, preventing tissue damage and maintaining immune tolerance. PD-L1 signalling counteracts T cell receptor stimulation, reducing T cell proliferation and cytotoxicity. Cancer cells exploit this pathway by aberrantly expressing PD-L1, disrupting immunosurveillance and facilitating immune evasion.

Immune checkpoint blockade (ICB) therapies targeting the PD-1/PD-L1 axis have become widespread and highly effective. However, a significant proportion of patients do not respond to these treatments, whilst some suffer adverse effects. Despite challenges due to the dynamic nature of PD-L1 expression within tumours and inconsistencies in diagnostic pathology methods, it has become evident that high PD-L1 expression is associated with better response to ICB in lung cancer ^3–6^. Thus, studying the cellular and genetic processes which regulate PD-L1 expression is an active research area aimed at improving response rates and patient outcomes.

Tumour PD-L1 expression involves critical transcriptional processes, including JAK-STAT-IRF1 signalling activated by interferon stimulation, Ras-MEK-ERK signalling triggered by receptor tyrosine kinase activation (e.g., EGFR signalling), and NF-κB activation ^7–9^. Epigenetic modifications at the CD274 locus also influence gene transcription ^10–13^. Post-transcriptionally, RNA-binding proteins and microRNAs can negatively affect PD-L1 mRNA stability and translation ^14–16^, while PD-L1 mRNA translation can be promoted through the integrated stress response ^17^. Post-translational modifications, such as glycosylation, phosphorylation, ubiquitination, acetylation, and palmitoylation are known to regulate PD-L1 protein stability and interactions ^18–23^.

Understanding how mutations in tumour suppressors and oncogenes trigger PD-L1 expression and facilitate immune evasion in distinct cancer types is crucial to improving response rates and patient outcomes. This knowledge can refine patient stratification for PD-1/PD-L1 ICB therapies, improving their predictability and efficacy ^24^.

LIMD1 encodes a 72 kDa protein with three LIM domains at its C-terminus, part of the Zyxin family ^25^. Recognized as a tumour suppressor, LIMD1 regulates hypoxia-inducible factor-1α (HIF-1α) and the Rb-E2F1 complex, influencing cell cycle progression and post-transcriptional gene regulation via microRNA-mediated gene silencing ^26–29^. LIMD1 also interacts with the Hippo signalling pathway, emphasising its role in diverse cellular processes ^30,31^. Loss of LIMD1 is common in NSCLC ^26^, but its implications for tumour-immune interaction remain unknown.

Here, we show that LIMD1 loss is clonal in a significant proportion of lung cancers. We investigate the impact of LIMD1 loss on tumour immunosurveillance, focusing on PD-L1 regulation. Via a novel scaffolding function with ARIH1, we find that LIMD1 is a critical regulator of PD-L1 protein turnover and restricts PD-L1 expression to maintain robust cancer immune surveillance. We report increased PD-L1 activity and attenuated T-cell-mediated target cell killing in the absence of LIMD1. Notably, tumours with low LIMD1 expression exhibit enhanced sensitivity to immune checkpoint inhibitors, underscoring the potential of LIMD1 as a biomarker for patient stratification in immunotherapy. Thus, LIMD1 is a crucial and previously unidentified regulator of PD-L1, highlighting therapeutic implications for lung cancer patients with LIMD1-deficient tumours.

## Results

### LIMD1 Loss is Clonal in Lung Cancer and is correlated with higher PD-L1 expression

Given that *LIMD1* is located at chr 3p21.3 which undergoes early loss in lung cancer ^32^, we initially investigated the TRAcking Cancer Evolution through therapy (Rx) (TRACERx) dataset of 421 NSCLC patients for LIMD1 loss of heterozygosity (LOH)^33^ ^34^. Multi-region sampling of tumours, along with both whole exome and RNA sequencing, permits the assessment of clonal gene mutation/ablation, gene expression signatures, and tumour evolution. We found that LIMD1 LOH is clonal in the majority of LUSC tumours (>80%), and 40% of LUAD, with subclonal loss in a further 20% of LUAD tumours (**Fig. 1A**). Based on these percentages, in the UK a conservative estimate is that approximately 20,000 patients diagnosed with NSCLC each year have LIMD1-deficiency/LOH (**Fig. 1B**). Consistent with the greater extent of LIMD1 LOH in LUSC, LIMD1 mRNA expression was significantly lower in LUSC compared to LUAD tumours. LIMD1 LOH correlated with the reduction in LIMD1 mRNA in LUAD. (**Fig. 1C**).

**Figure 1.**
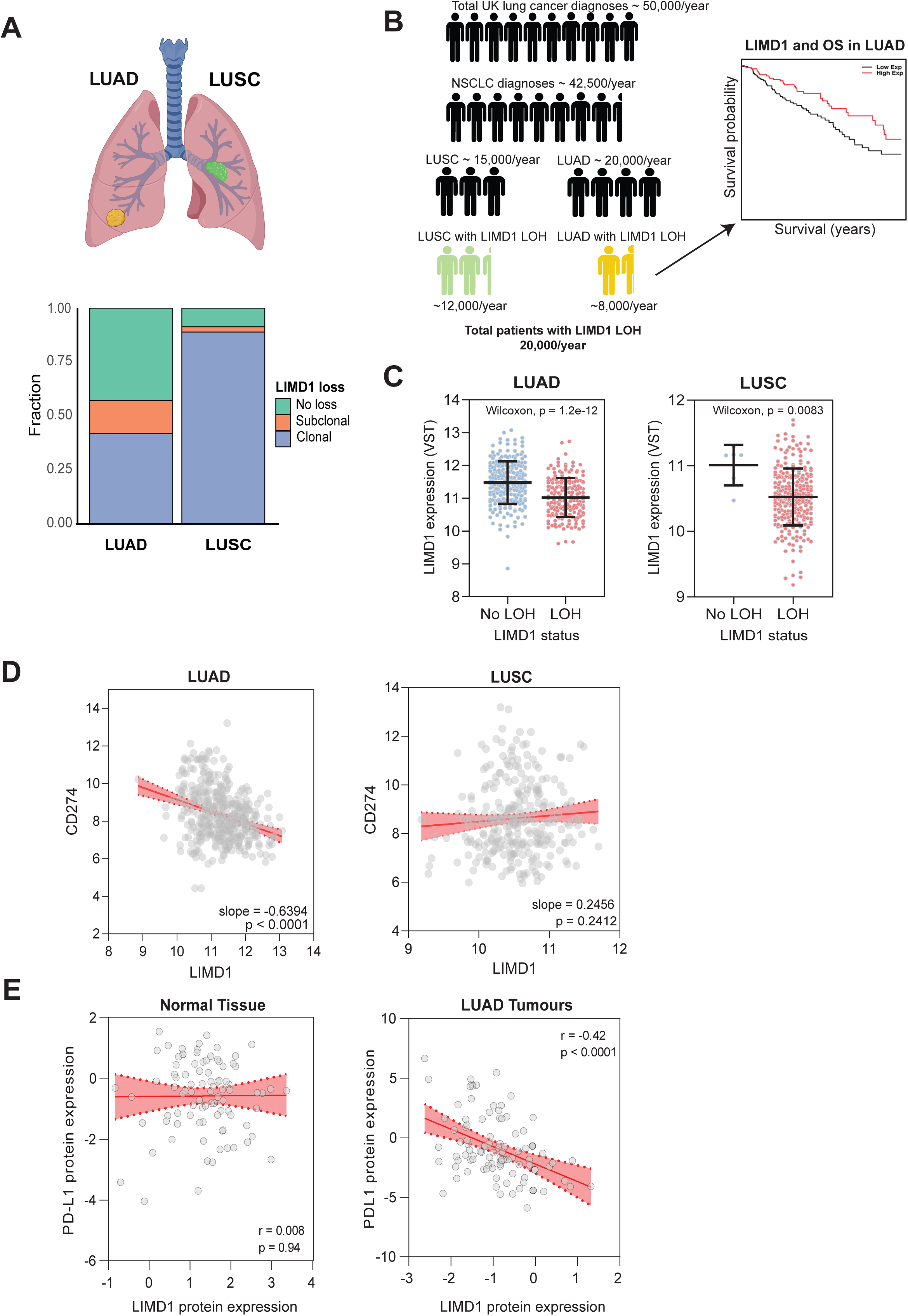
Clonal LIMD1 LOH is prevalent in both LUSC and LUAD and correlated with PD-L1 expression in LUAD. (**A**) Proportion of samples in the TRACERx cohort exhibiting clonal or subclonal LOH for LIMD1 in LUAD and LUSC. (**B**) Estimates of numbers of lung cancer patients in the UK affected by LIMD1 LOH. One ‘stick man’ figure is equivalent to 5,000 patients. (**C**) LIMD1 mRNA expression in LUAD and LUSC patient samples exhibiting LOH or no LOH for LIMD1. Line shows median LIMD1 expression ± the interquartile range. P values were calculated according to the Wilcoxon signed-rank test. (**D**) Correlation analysis of LIMD1 and PD-L1 mRNA expression from patients from the TRACERx-421 cohort. (**E**) Proteomic analysis quantifying LIMD1 levels and PD-L1 levels in LUAD and matched healthy adjacent tissue (n=100). Data obtained from the Clinical Proteomic Tumor Analysis Consortium’s (CPTAC) dataset of lung adenocarcinomas. For D & E, lines of best fit were determined with linear regression analysis and red shaded areas denote the 95% confidence intervals.

The clonal loss of LIMD1 suggests a fundamental role in lung cancer pathogenesis and tumour cell evolution. Given the established association between tumour suppressor loss and immune checkpoint regulation ^35,36^, we investigated the relationship between LIMD1 deficiency and PD-L1 expression across NSCLC subtypes. Our analysis focused on both mRNA and protein levels within available datasets to provide a comprehensive view of this potential regulatory interaction. Within the TRACERx 421 dataset, we observed a significant negative correlation between LIMD1 mRNA expression and CD274 mRNA in LUAD (**Fig. 1D**). In LUSC, however, this correlation was not apparent. Analysis of the Clinical Proteomic Tumour Analysis Consortium (CPTAC) dataset ^37^ also revealed a significant inverse correlation between LIMD1 and PD-L1 protein in LUAD tumours, a trend that was not present in normal adjacent tissue (**Fig. 1E**). No such correlation was seen for LUSC in this dataset (data not shown). Thus, across multiple cohorts of LUAD patients, we find a significant LIMD1 LOH and an inverse association with PD-L1 expression.

Our previous work has defined LIMD1 loss as a driver of dysregulation of the cellular response to hypoxia ^26,38^. In keeping with this, we found in the TRACERx 421 cohort that lower expression of LIMD1 in LUAD and LUSC samples correlated with a higher hypoxia metagene signature, as defined by Buffa et al., ^39^ (**Supp Fig. 1A**). In the same cohort of LUAD and LUSC patients, we visualised CD274 expression against the Buffa et al., hypoxia metagene signature (**Supp Fig 1B**). We observed a significant positive correlation between the two variables in LUAD, whereas no such correlation was found in LUSC. These findings indicate that LIMD1 loss may contribute to immune evasion in LUAD through a mechanism linked to hypoxia-related pathways.,

### LIMD1 Negatively Regulates PD-L1 Expression

The observed correlation between LIMD1 and PD-L1 levels in patient samples led us to investigate the molecular mechanism underpinning this. We first examined whether this relationship is present in cancer cell lines *in vitro*, which would allow us to dissect the regulatory pathway more effectively. LIMD1 depletion by RNA-interference (RNAi) with short-hairpin RNA (shRNA) upregulated PD-L1 protein in an range of cancer cell types, including H1299 and A549 LUAD, and RCC48 renal carcinoma cells. Re-expression of an RNAi-resistant LIMD1 expression cassette rescued control of PD-L1 expression and reduced it to levels comparable to that of non-targeting shRNA controls (**Fig. 2A,B**), demonstrating that LIMD1 is both necessary and sufficient for PD-L1 regulation at the protein level in these cells.

**Figure 2.**
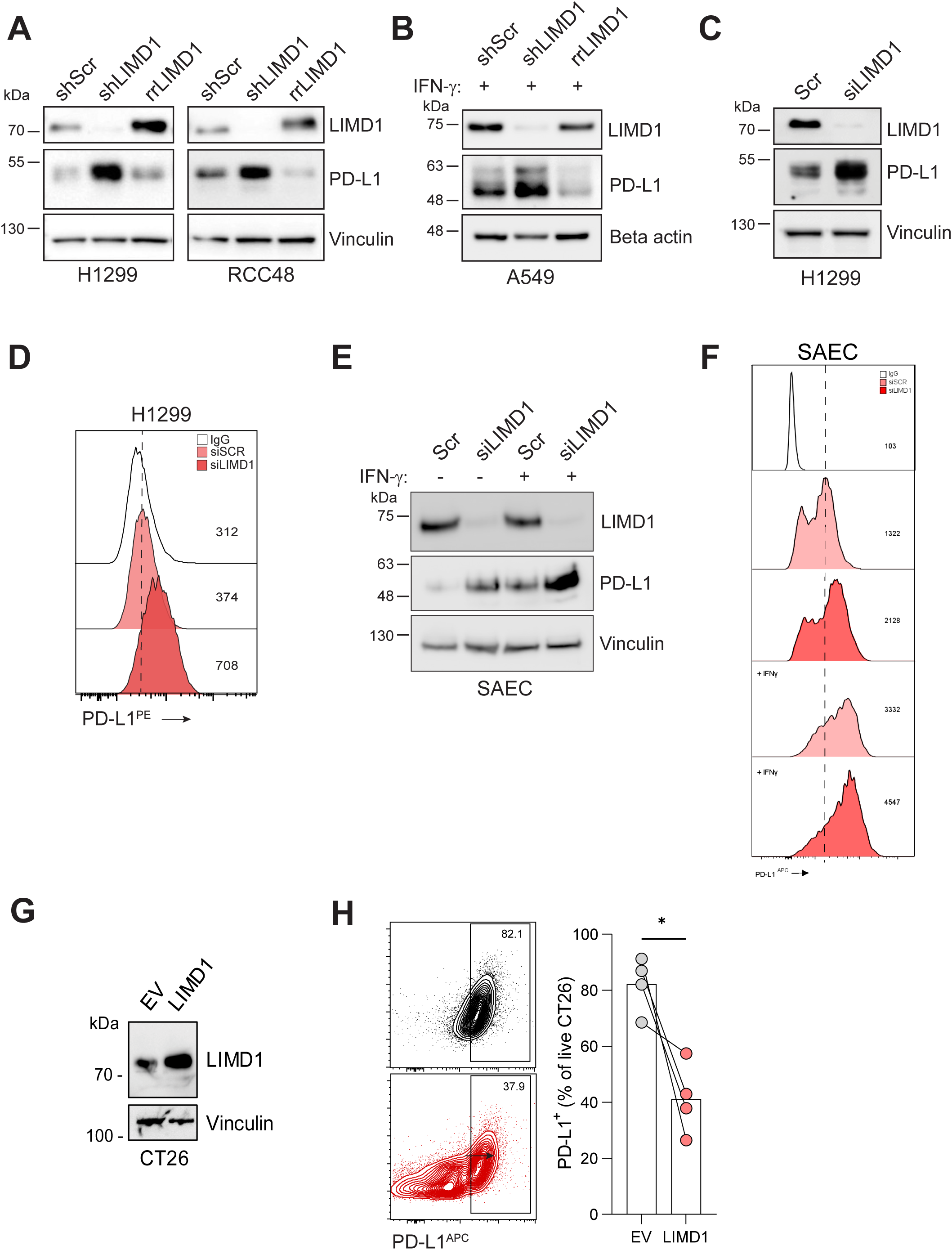
LIMD1 regulates PD-L1 expression in multiple primary and transformed lung cell types. (**A & B**) Immunoblots of PD-L1 protein in H1299 LUAD cells, RCC48 renal carcinoma cells and A549 LUAD cells transduced with shRNA for LIMD1 and RNAi-resistant LIMD1 lentiviral vectors. A549 cells were treated with 2 ng/mL IFNγ overnight. Vinculin was probed as a loading control. (**C & D**) Immunoblot and flow cytometric analysis showing PD-L1 expression in H1299 cells following LIMD1 silencing with siRNA (20nM). (**E**) Immunoblots for PD-L1 and LIMD1 in SAEC whole cell lysate upon LIMD1 knockdown with siRNA and treatment with 0.5 ng/mL IFNγ overnight. Vinculin was probed as a loading control. (**F**) Flow cytometric analysis of surface PD-L1 in SAEC transfected with siScr or siLIMD1, +/- treatment with 0.5 ng/mL IFNγ overnight. Values indicate median fluorescence intensity. (**G**) Immunoblot showing stable LIMD1 over-expression in CT26 colorectal carcinoma cells. (**H**) Flow cytometric analysis of surface PD-L1 in CT26. Data represent three independent replicates (*p < 0.05, paired student’s t-test)

Similarly, LIMD1 knockdown in H1299, A549, H3122, and H358M LUAD models increased PD-L1 protein, particularly following interferon-gamma (IFNγ) treatment (**Fig. 2 B-D and Supp. Fig. 2A**). In primary human small airway epithelial cells (SAEC), we again observed upregulation of PD-L1 protein upon LIMD1 knockdown (**Fig. 2E,F**). Furthermore, LIMD1-dependent control of PD-L1 expression was also observed in murine cells, including CT26 colorectal carcinoma and MCA-205 fibrosarcoma where Limd1 was overexpressed (**Fig. 2 G,H and Supp. Fig. 2B,C**). Interestingly, depletion of Ajuba, a paralogous protein to LIMD1 that is frequently inactivated in pleural mesothelioma models^40^, also led to dramatic increases in PD-L1 expression (**Supp. Fig. 2D**). This finding implies that members of the Zyxin family may share functional roles in regulating PD-L1 expression.

To further probe the LIMD1-PD-L1 regulatory relationship, A549 LIMD1^-/-^ cells were generated using CRISPR-Cas9 editing. Two LIMD1 knockout clones (C5 and C10) were studied alongside a Cas9-expressing LIMD1^+/+^ control cell line. In keeping with RNAi models, A549 LIMD1^-/-^ had markedly elevated PD-L1 expression after inflammatory challenge with conditioned-medium from anti-CD3 activated PBMCs, likely due to a combination of IFNγ and TNFα signalling (**Fig. 3A-C**). Indeed, A549 LIMD1^-/-^ exhibited significantly higher IFNy-induced PD-L1 expression than LIMD1^+/+^ cells (**Fig. 3D and Supp Fig 3A**). Despite the striking difference in PD-L1 protein expression, PD-L1 mRNA levels were similar between knockout and control cells (**Fig. 3E Supp. Fig 3B**), a trend also observed in wild-type A549 following transient LIMD1 depletion with siRNA (**Fig. 3F,G**). These data indicate that in A549, LIMD1 loss does not enhance the canonical IFNγ-JAK-STAT signalling pathway, a known driver of PD-L1 transcription.

**Figure 3.**
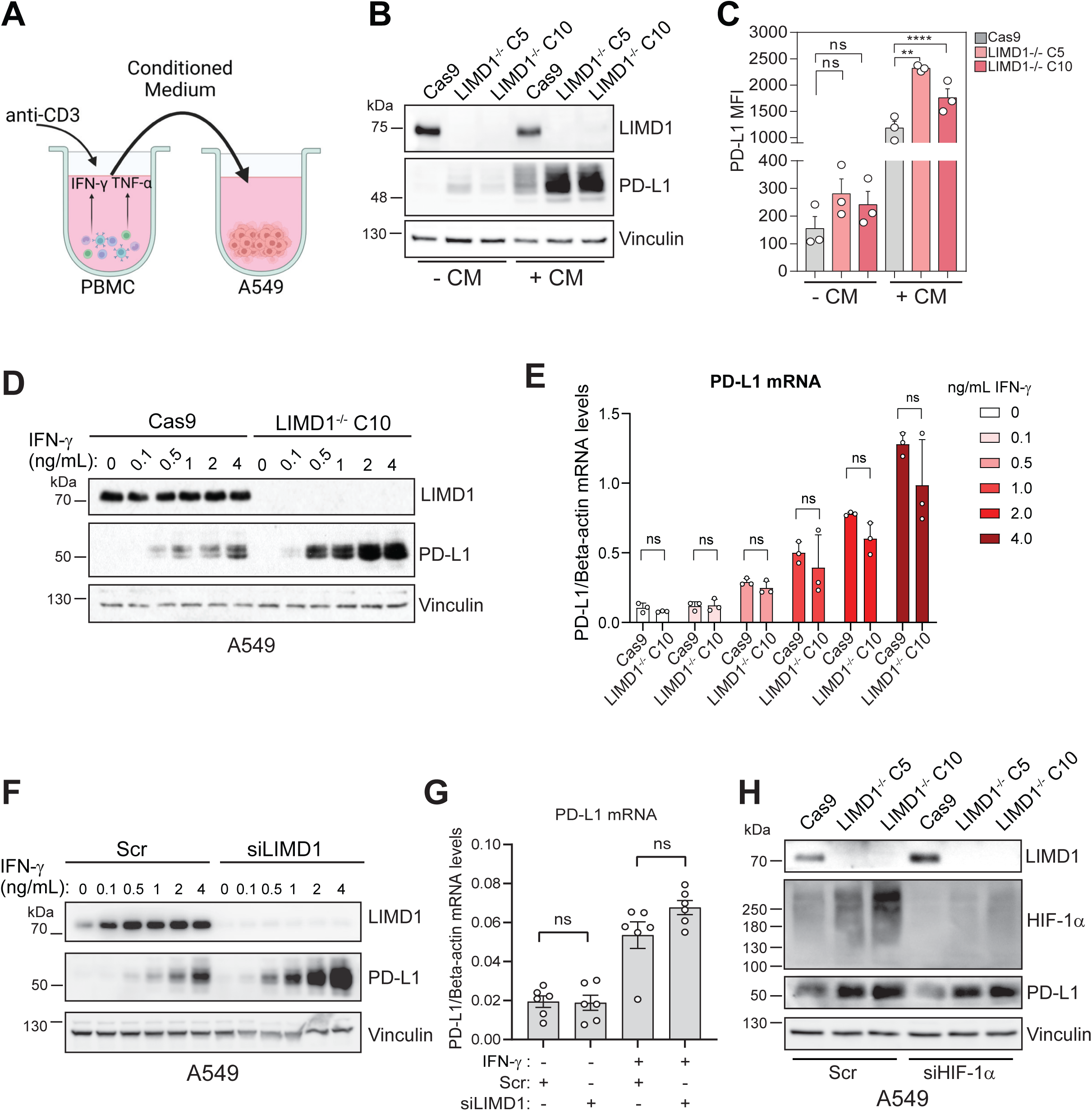
Loss of LIMD1 in A549 increases PD-L1 expression at the protein level. (**A**) Schematic showing collection of conditioned medium (CM) from PBMCs activated with anti-CD3 (5 µg/mL, 3 days) for induction of PD-L1 expression in A549. Immunoblot (**B**) and flow cytometric analysis (**C**) of Cas9 control or LIMD1-ablated A549 treated with PBMC-conditioned medium for 24 h. The histogram shows PD-L1 surface expression in each group. Three biological replicates (mean, ±s.e.m). ****p <0.0001, **p <0.01, *p <0.05 from one-way ANOVA with Sidak’s multiple comparisons test. Analysis of PD-L1 protein by immunoblot (**D**) and mRNA levels by qPCR (**E**) in LIMD1-knockout (LIMD1^-/-^ C10) and control (Cas9) A549 following stimulation with various concentrations of recombinant IFNγ for 24 hours. The graph shows mean ±s.e.m. CD274 mRNA levels normalised to beta-actin mRNA levels of three independent experiments. n.s. = non-significant according to one-way ANOVA with Sidak’s multiple comparisons test. (**F**) PD-L1 immunoblot in WT A549 transfected with LIMD1 siRNA or control siRNA (Scr) and treated with various concentrations of recombinant IFNγ for 24 hours. (**G**) qPCR of PD-L1 mRNA in WT A549 transfected with Scr or LIMD1 siRNA and treated with Vehicle or 1 ng/mL recombinant IFNγ for 24 hours. Data shows mean ± s.e.m PD-L1 mRNA normalised to beta-actin mRNA for six independent experiments. N.s. = non-significant according to the Mann-Whitney test. (**H**) A549 LIMD1^+/+^ control and LIMD^-/-^ clones were transfected with non-targeting siRNA (Scr) or siHIF-1ɑ. Immunoblotting was performed for HIF-1ɑ, PD-L1 and LIMD1. Vinculin was probed as a loading control.

Guided by the strong positive correlation between CD274 mRNA and hypoxic gene expression signature in LUAD samples (**Supp Fig 1B**), we next examined if the increased PD-L1 expression in A549 LIMD1^-/-^ was dependent on HIF-1ɑ. As expected from previous findings^26^, HIF-1ɑ displayed increased levels in LIMD1^-/-^ cells. However, since HIF-1ɑ silencing did not alter PD-L1 protein levels (**Fig. 3H**), it is unlikely that it contributes to the PD-L1 enrichment observed in LIMD1^-/-^ cells. Furthermore, despite the HIF-1ɑ enrichment classically seen in LIMD1-deficient cells, PD-L1 mRNA was consistent between LIMD1^+/+^ and LIMD1^-/-^ isogenic A549 (**Fig. 3E**), thus HIF-1ɑ transcriptional activity at the CD274 locus is not responsible for the PD-L1 deregulation observed.

Overall, loss of LIMD1 increased PD-L1 expression in multiple cell types and different mammalian species, uncovering a novel evolutionarily-conserved cancer cell-intrinsic mechanism controlling immune checkpoint expression.

### LIMD1 Loss does not affect PD-L1 expression post-transcriptionally

Given the observed discrepancy between PD-L1 protein and mRNA levels in LIMD1-deficient A549, we investigated post-transcriptional mechanisms that may regulate PD-L1 expression. After excluding HIF-driven pathways, we focused on the known interaction of LIMD1 with components of the microRNA-induced silencing complex (miRISC), such as Argonaute (AGO) and TNRC6 proteins, which are involved in post-transcriptional gene silencing. We hypothesised that the LIMD1 loss may affect the regulation of the PD-L1 3’UTR, a region of the mRNA targeted by miRNA-mediated silencing ^15,27,28^. However, depletion of LIMD1 in A549 did not alter the expression of a Renilla luciferase reporter gene containing the PD-L1 3’UTR (**Supp. Fig. 4A**). In contrast, knockdown of Ajuba and AGO1 led to de-repression of this reporter gene. These results suggest that the role of LIMD1 in miRNA-mediated silencing is not responsible for the increased PD-L1 protein expression observed following LIMD1 loss, but related family-member proteins exhibit the capacity to post-transcriptionally regulate PD-L1 at the 3’UTR. This suggested that the increased PD-L1 protein levels in LIMD1-deficient cells may be due to post-translational mechanisms.

### Impaired PD-L1 Degradation in LIMD1-Deficient Cells

Testing whether loss of LIMD1 affected PD-L1 protein stability, we used cycloheximide (CHX) treatment and found significant stabilisation of PD-L1 in A549 LIMD1^-/-^ cells (**Fig. 4A,B**). We next investigated specific degradation pathways that may be involved, namely lysosomal and proteasomal degradation mechanisms. To test for altered lysosomal degradation, we treated A549 with chloroquine (CQ), a potent inhibitor of lysosomal acidification. As expected, LC3B levels increased with CQ treatment, however in both control and LIMD1^-/-^ cells, PD-L1 decreased to a similar extent (**Supp. Fig. 4B**).

**Figure 4.**
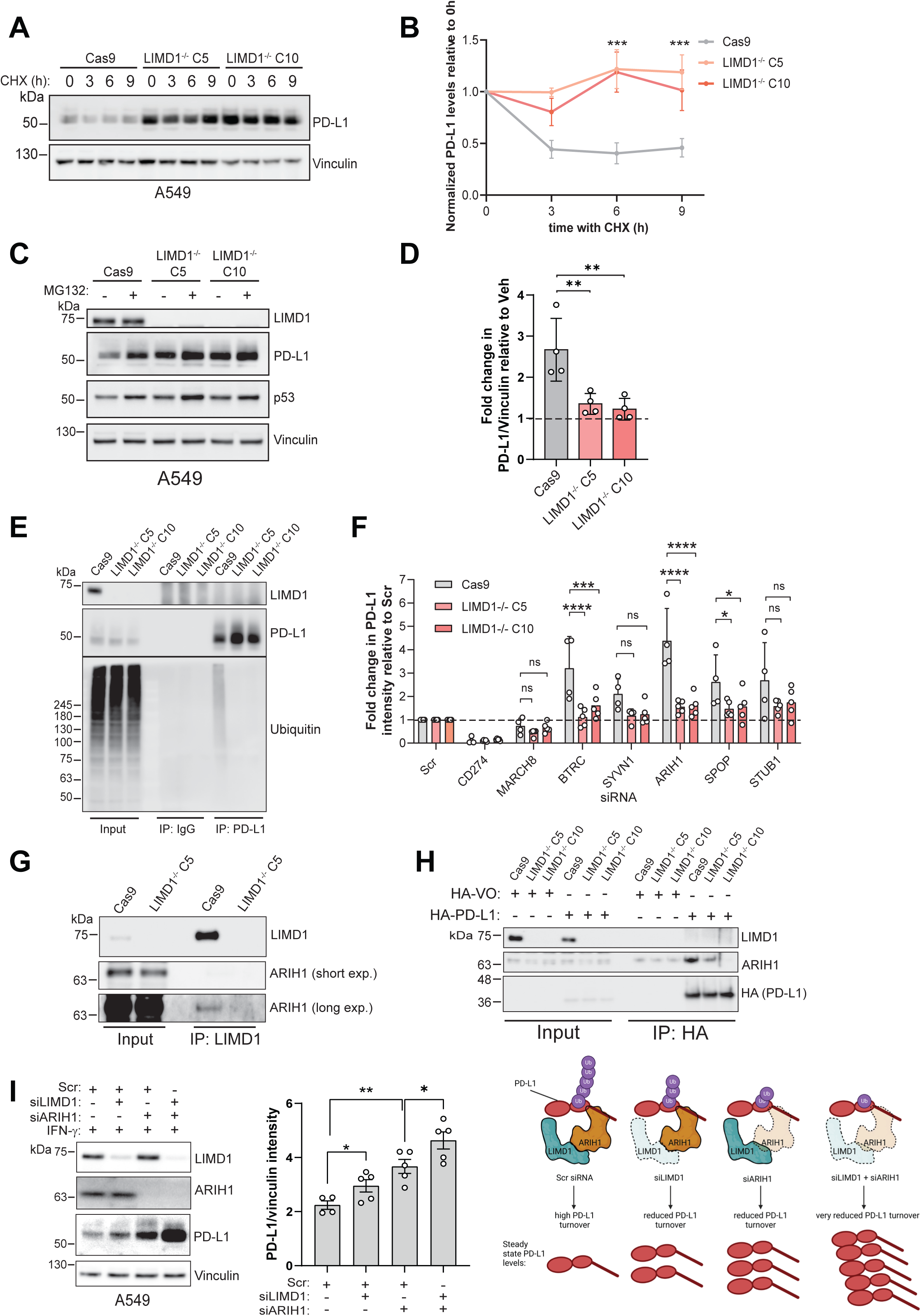
PD-L1 upregulation following LIMD1 loss is partly driven by increased PD-L1 protein stability and reduced proteasomal degradation. **(A)** Immunoblot analysis of PD-L1 in A549 Cas9 and LIMD1^-/-^ cells treated with 40 μg/mL cycloheximide for the indicated time points. Vinculin was probed as a loading control. (**B**) Graph shows mean fold change in PD-L1 level following CHX treatment determined by densitometry relative to the 0h time point normalised to vinculin levels for three independent experiments ± s.e.m. **(C)** Immunoblot of cells treated with 10 μM MG132 or vehicle for 24h. (**D**) Graph shows mean fold change in PD-L1 relative to the vehicle upon 24h MG132 treatment for four independent experiments ± s.e.m. Vinculin was probed as a loading control. **(E)** A549 Cas9 and LIMD1^-/-^ C10 cells were treated with IFNγ for 24h and with 10 µM MG132 for 6h. PD-L1 was then immunoprecipitated from whole cell lysate and elutions and input samples were immunoblotted for ubiquitin. Control IPs were performed with rabbit IgG. **(F)** Immunoblot densitometry of fold change in PD-L1 relative to Scr siRNA transfected A549 Cas9 or LIMD1^-/-^ C5 and C10 cells upon siRNA depletion of various PD-L1 regulators or CD274 itself. PD-L1 density was normalised to corresponding vinculin loading control. Data shows mean fold change for at least four independent experiments ± s.e.m. **(G)** Immunoblots of endogenous co-immunoprecipitation of ARIH1 with LIMD1 in A549 Cas9 and C5 LIMD1^-/-^ cells. **(H)** Co-immunoprecipitation of HA-PD-L1 in A549 Cas9 and LIMD1^-/-^ cells with ARIH1. **(I)** PD-L1 immunoblot following LIMD1 or ARIH1 or combined depletion with siRNA in WT A549 cells treated with 2 ng/mL IFNγ. Vinculin was probed as a loading control. Graph shows mean densitometry for PD-L1 normalised to vinculin intensity for at least four independent experiments. **p <0.01, *p <0.05 for unpaired t-test. Schematic illustrates the effect of depleting one or both parts of a complex on PD-L1 levels. (**J**) Schematic of LIMD1’s regulation of ARIH1-dependent turnover of PD-L1 in lung cancer cells.

Assessment of proteasome function using the inhibitor MG132 revealed significant PD-L1 enrichment in A549 LIMD1^+/+^ only, suggesting impaired degradation in LIMD1-deficient cells (**Fig. 4C,D**). This was supported by immunoprecipitation data showing reduced poly-ubiquitination of PD-L1 in A549 LIMD1^-/-^ (**Fig. 4E**), indicating that the high PD-L1 expression level characteristic of LIMD1-deficient cells is, at least partially, a result of impaired ubiquitination and PD-L1 proteasomal turnover.

### ARIH1 Mediates PD-L1 Ubiquitination in a LIMD1-Dependent Manner

Recent studies have identified several regulators involved in the ubiquitination of PD-L1 (reviewed in ^41^). To identify E3 ubiquitin ligases which may differentially regulate PD-L1 depending on LIMD1 status, we used a candidate approach to silence known regulators and measure PD-L1 level in A549 cells. (**Fig. 4F** and **Supp. Fig. 4C**). Interestingly, we observed a marked and selective relative increase in PD-L1 in A549 LIMD1^+/+^ over LIMD1^-/-^ clones following depletion of the E3 ligases BTRC (βTrCP) or ARIH1 (**Fig. 4F**), which have been shown to mediate PD-L1 ubiquitination by others ^19,42^. Conversely, depletion of other E3 ligases, including MARCH8, SYVN1, and STUB1, did not modulate PD-L1 differentially between LIMD1^-/-^ and control cells.

Given the precedent of LIMD1 interacting with E3 ubiquitin ligases (e.g., VHL ^38^), we confirmed that LIMD1 directly interacts with ARIH1 by endogenous co-immunoprecipitation in A549 (**Fig. 4G**). Closer examination of molecular interactions revealed that co-immunoprecipitation of HA-PD-L1 with ARIH1 was reduced in A549 LIMD1^-/-^ cells (**Fig. 4H**). This reveals a previously unrecognised role for LIMD1 in facilitating the interaction between PD-L1 and ARIH1, suggesting that LIMD1 may generally enhance the functionality of E3 ubiquitin ligases in regulating protein turnover. With independent siRNA, we found that co-depletion of LIMD1 and ARIH1 in WT A549 significantly augmented the deregulation of PD-L1 (**Fig. 4I**), whereas depleting them individually led to moderate increases in PD-L1. This experiment indicates that LIMD1 and ARIH1 interact to regulate PD-L1 turnover, with their combined activity promoting highly efficient PD-L1 ubiquitination and degradation. In the absence of LIMD1, ARIH1-dependent PD-L1 ubiquitination and degradation is impaired and so PD-L1 protein accumulates.

### LIMD1 Loss Modulates Tumour-Immune Cell Interactions

Building on our mechanistic insights into the regulation of PD-L1 by LIMD1, we sought to determine the functional relevance of this regulatory axis within the context of immune surveillance and tumour-immune interactions. To this end, we conducted a series of *in vitro* experiments designed to evaluate how LIMD1 loss influences human CD8^+^ T cell activation and tumour sensitivity to immune checkpoint inhibition. These *in vitro* assays allowed us to directly explore human-specific cellular mechanisms.

To validate the functional effect of PD-L1 upregulation on tumour-immune cell interactions, we examined human co-cultures of peripheral blood mononuclear cells (PBMCs) with LIMD1-deficient target cancer cells. In keeping with A549 LUAD cells, the LUAD model H1299 exhibited elevated surface PD-L1 following LIMD1 deletion (**Fig. 5A**) and were significantly more suppressive than control cells in inhibiting activated CD8^+^ T cell proliferation (**Fig. 5B,C**). Measurement of CD25 expression in a similar allogeneic culture model also revealed reduced CD4^+^ and CD8^+^ activation in A549 LIMD1^-/-^ co-cultures (**Fig. 5D**). Interestingly, treatment with the PD-L1 inhibitor atezolizumab increased CD25 expression in CD8^+^ T cells within LIMD1-deficient cultures to a greater extent than in those with LIMD1^+/+^ cells, indicating that tumour cell LIMD1 loss sensitised T cells to immune checkpoint inhibition *in vitro*.

**Figure 5.**
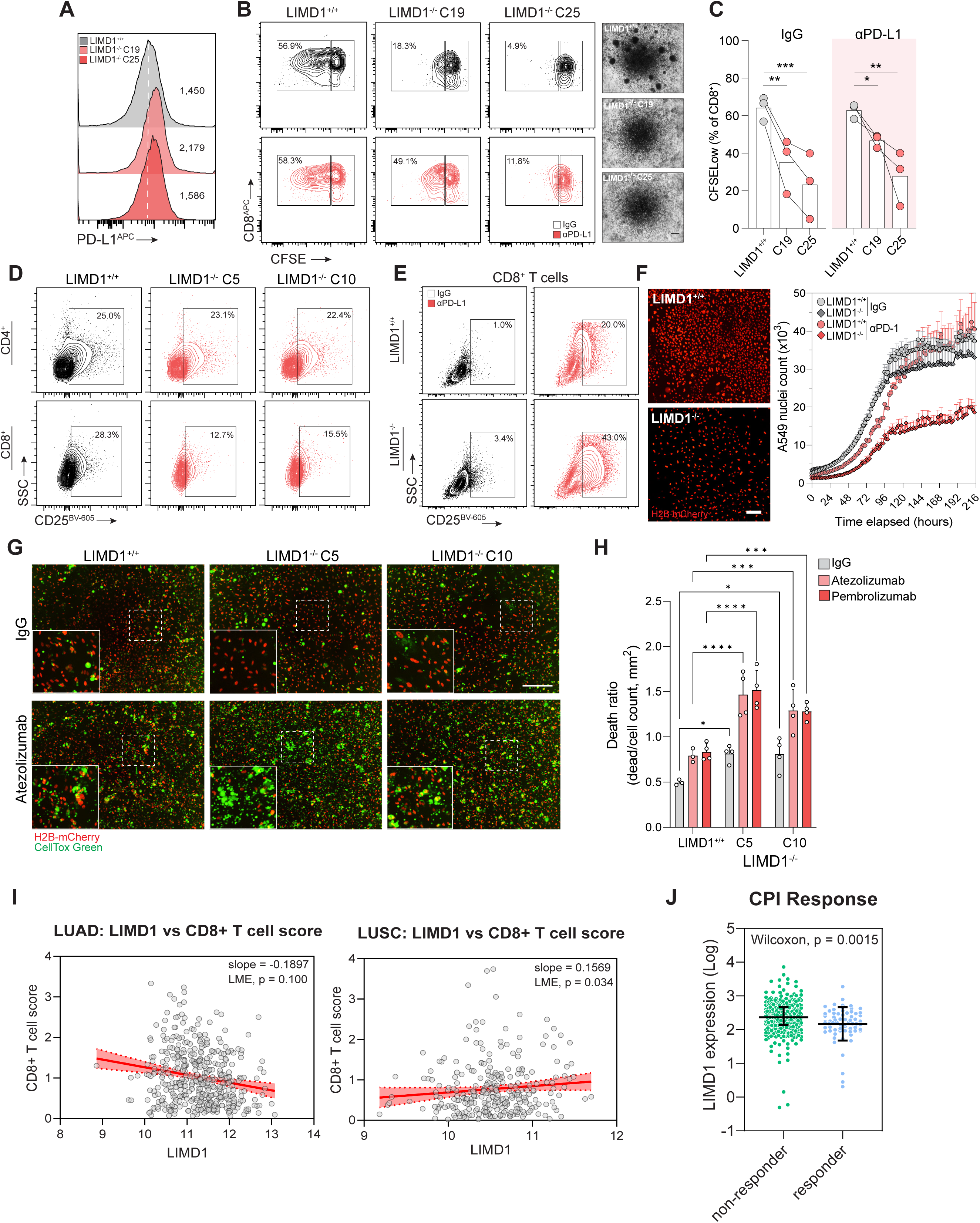
LIMD1-deficient cells suppress T-cell activation and are sensitive to PD-L1 blockade *in vitro*. (**A**) Surface PD-L1 expression in H1299 cells measured by flow cytometry. Values indicate median fluorescence intensity. (**B**) CD8^+^ T cell proliferation in co-culture with H1299 target cells with or without anti-PD-L1 (10 µg/mL). Phase-contrast images show representative cultures after 5 days (scale = 100µm). (**C**) CD8^+^ T cells proliferation within H1299 co-cultures. ***p <0.001, **p <0.01, *p <0.05 for two-way ANOVA with Holm-Sidak correction. Data represent PBMC from n=3 independent healthy donors. **(D)** Flow cytometric measurement of CD25 expression in T cells following 5-day co-culture with A549 (untreated). (**E**) Measurement of CD8^+^ T cell proliferation within PBMC-A549 co-cultures following ICB with atezolizumab (10 µg/mL). (**F**) Cell proliferation curves showing A549 growth within PBMC co-cultures following ICB with atezolizumab. Representative end-point images show A549 nuclei density at 200 hours (scale = 300 μm). (**G-H**) Live-cell microscopy measurement of cytotoxicity within A549-PBMC 2D co-cultures treated with atezolizumab or IgG control (10ug/mL). Representative end-point images show nuclei density and cell death signal at 150 hours (scale bar = 150μm). Data are mean ± sem, n=4 (****p < 0.0001, ***p < 0.001, *p <0.05, two-way ANOVA with Holm-Sidak correction). (**I**) LIMD1 mRNA expression vs CD8+ T cell score in LUAD and LUSC tumours. Red line is the line of best fit according to simple linear regression with shaded areas representing 95% confidence intervals. P value was calculated using a linear mixed-effects model. (**J**) LIMD1 mRNA expression (log-transformed) in NSCLC patients, taken from the Patil et al., and Banchereau et al. cohorts, stratified by response to immune checkpoint blockade therapy.

To assess how LIMD1 loss affects tumour cell fitness within co-cultures, we cultured A549 fluorescent reporter cells with activated PBMCs. Although both cell types exhibited comparable growth rates in IgG-treated cultures, we observed a significant and selective reduction in A549 LIMD1^-/-^ proliferation following treatment with the PD-1 inhibitor pembrolizumab (**Fig. 5E**). Incorporation of cytotoxicity dyes revealed significantly more apoptosis in atezolizumab-treated co-cultures in the absence of LIMD1 (**Fig. 5F,G**). A similar pattern was observed with 3D co-cultures of tumour cell spheroids and activated PBMCs, with LIMD1^-/-^ spheroids exhibiting marked cytotoxicity relative to LIMD1^+/+^ controls following treatment with both PD-1 and PD-L1 inhibitors (**Supp. Fig. 5A,B**). Given the functional effects of LIMD1 loss on T cell activation *in vitro,* we reanalysed the TRACERX 421 patient data for intratumoural T cell signatures. Interestingly, we found a correlation between LIMD1 expression and CD8+ T cell in tumour samples from LUSC – low LIMD1 expression associated with lower CD8+ T cell score (**Fig. 5I**). Of note we did not see such an association in LUAD.

### Low LIMD1 Expression Correlates with Response to Immune Checkpoint Inhibitors in NSCLC

To investigate the relationship between LIMD1 expression and response to immune checkpoint inhibitors (ICIs) in non-small cell lung cancer (NSCLC), we analysed LIMD1 expression levels in relation to clinical response, using data from two cohorts ^43,44^. Our analysis revealed that patients with low LIMD1 expression were significantly more likely to respond to ICIs, as indicated by the correlation between decreased LIMD1 expression and response status (Wilcoxon p = 0.0015; **Fig. 5J**). Stratifying NSCLC patients based on their response status (responder vs. non-responder) showed that responders exhibited lower LIMD1 expression compared to non-responders (**Fig. 5J**). These findings suggest that low LIMD1 expression may serve as a predictive biomarker for ICI efficacy in NSCLC.

In summary, these results underscore that LIMD1 loss is a clonal event in a significant proportion of NSCLC cases. We demonstrate that LIMD1 regulates PD-L1 at the post-translational level, with its loss leading to impaired PD-L1 degradation via the ARIH1 E3 ubiquitin ligase complex. This impairment increases PD-L1 protein stability, thereby promoting immune evasion. Additionally, we reveal that low LIMD1 expression correlates with a positive response to immune checkpoint inhibitors (ICIs) in NSCLC, suggesting that LIMD1 may serve as a predictive biomarker for ICI efficacy. Importantly, these regulatory effects of LIMD1 on PD-L1 stability are conserved across multiple cancer cell types, including lung and colorectal carcinoma, as well as mouse cell lines.

## Discussion

Immune evasion via PD-L1 upregulation is a hallmark of tumour progression and a critical barrier to effective immunotherapy in non-small cell lung cancer (NSCLC). While transcriptional regulation of PD-L1 through pathways like JAK-STAT^45^ and hypoxia-induced HIF-1α signalling^46^ has been well characterized, the role of post-translational mechanisms in PD-L1 stabilization remains less understood. Here, we identify LIMD1, a tumour suppressor gene frequently lost in NSCLC, as a novel regulator of PD-L1 stability through its interaction with the E3 ubiquitin ligase ARIH1. LIMD1 loss disrupts ARIH1-mediated ubiquitination and proteasomal degradation of PD-L1, leading to its stabilization and enhanced immune evasion.

### LIMD1 Loss as an Early and Clonal Event in Tumour Evolution

Our data demonstrate that LIMD1 loss is a frequent and clonal event in NSCLC, with >80% of lung squamous cell carcinoma (LUSC) and ∼40% of lung adenocarcinoma (LUAD) cases exhibiting loss of heterozygosity (LOH). This clonal nature highlights LIMD1 loss as an early driver of tumour evolution, distinguishing it from other PD-L1 regulators that are often upregulated later in tumour progression. Previous studies have linked truncal mutations in tumour suppressors like LKB1, PTEN, and TP53 to PD-L1 regulation via transcriptional pathways ^24,47,48^. In contrast, our findings reveal a distinct post-translational mechanism, positioning LIMD1 as a critical sentinel for PD-L1 turnover during early oncogenesis.This distinction is particularly relevant in LUAD, where we observe a significant inverse correlation between LIMD1 expression and PD-L1 levels at both mRNA and protein levels. The absence of a similar correlation in LUSC may reflect tumour-specific differences in immune escape mechanisms, with LUSC potentially relying on alternative pathways. These findings underscore the importance of integrating tumour-specific genetic contexts when considering immune checkpoint regulation.

### Mechanistic Insights into LIMD1-ARIH1 Regulation of PD-L1

We have shown that LIMD1 loss in A549 cells increases PD-L1 protein stability. Unlike previous reports of CQ-induced PD-L1 enrichment in gastric cancer cells ^49^, our findings indicate that PD-L1 does not undergo significant lysosomal degradation in A549. Instead, we found that proteasomal degradation of PD-L1 is reduced with LIMD1 loss. In keeping with this, we found greater levels of poly-ubiquitinated PD-L1 in LIMD1 expressing A549 cells, suggesting that LIMD1 loss impairs ubiquitin-dependent proteasomal degradation of PD-L1. Previous work on LIMD1 has shown that its LIM domains mediate protein-protein interactions. During the regulation of HIF-1α, LIMD1 interacts via its LIM-domains with VHL, the E3 ligase which ubiquitinates HIF-1α ^38^. We now show a specific interaction between LIMD1 and the E3 ligase ARIH1, and this finding is consistent with recent studies showing that ARIH1 regulates PD-L1 stability through the ubiquitin-proteasome system ^42^. While Liu et al. highlighted ARIH1’s role in regulating PD-L1 following chemotherapy^50^, our work demonstrates that LIMD1 is essential for ARIH1 function under basal conditions in the absence of inflammatory stimuli or chemotherapy. Specifically, LIMD1 interacts with the ARIH1 to facilitate PD-L1 ubiquitination and subsequent proteasomal degradation. When LIMD1 is absent, the ARIH1-PD-L1 interaction is disrupted, resulting in reduced poly-ubiquitination and impaired degradation of PD-L1, thereby increasing its steady-state levels. This novel regulatory mechanism highlights LIMD1 as a critical modulator of PD-L1 expression and underscores its potential role in tumour immune evasion.

Interestingly, depletion of ARIH1 alone resulted in greater PD-L1 stabilization compared to LIMD1 loss, suggesting that ARIH1 may function partially independently of LIMD1. This raises the possibility that ARIH1 interacts with other regulators or substrates in the absence of LIMD1, providing redundancy in PD-L1 turnover pathways. Nevertheless, our data highlight LIMD1 as a critical modulator of ARIH1-dependent PD-L1 regulation, adding a new layer of complexity to immune checkpoint biology.

As PD-L1/*CD274* transcription is exquisitely sensitive to IFNγ signalling (and other inflammatory stimuli), in non-inflammatory conditions LIMD1-dependent PD-L1 protein turnover may be critical for minimising PD-L1 protein levels that may arise from low level transcription. In this context, LIMD1 loss and consequent impairment of PD-L1 ubiquitination and/or proteasomal degradation may lead to a steady accumulation in PD-L1 over time. With IFNγ treatment, we and others have shown transcription of *CD274* increases dramatically. Notably, in A549 cells we found LIMD1 loss did not affect *CD274* transcript levels and yet protein levels of PD-L1 were significantly higher in LIMD1^-/-^ cells. Thus, loss of LIMD1 in both non-inflammatory and inflammatory contexts may upregulate PD-L1 protein levels, which facilitate tumour immune evasion. Furthermore, while ARIH1 seems to be the primary E3 ligase regulating PD-L1 in A549 cells, it is possible that LIMD1 interacts with other E3 ligases in different tissue types or under different stress conditions, a hypothesis that warrants further investigation.

### LIMD1 and Immune Checkpoint Blockade Response

Clinically, PD-L1 expression has long been considered the primary biomarker for predicting response to immune checkpoint inhibitors (ICIs), such as pembrolizumab and nivolumab, particularly in NSCLC. However, while PD-L1 testing remains a widely used predictor in clinical practice, its utility is limited due to the dynamic and heterogeneous nature of PD-L1 expression within tumours, leading to variability in response rates. Emerging evidence suggests that other biomarkers, such as tumour mutational burden (TMB), tumour-infiltrating lymphocytes (TILs), and specific genomic signatures, may provide more robust or complementary predictive value for ICI response ^51,52^. Despite these advancements, PD-L1 expression remains a key factor in clinical decision-making, and our findings that LIMD1 loss drives PD-L1 stabilisation (Fig. 5I) offers important insights into immune checkpoint regulation, particularly in a subset of NSCLC patients. That said, more work is needed to understand this biology completely, as the range of LIMD1 expression in non-responders significantly overlaps with that of responders, preventing LIMD1 loss alone from being classified as a definitive biomarker. This overlap may reflect the multifactorial nature of ICI response, where PD-L1 stabilisation alone is insufficient to predict outcomes. Factors such as tumour mutational burden (TMB), CD8^+^ T cell infiltration, and the broader tumour immune microenvironment likely interact with LIMD1 loss to influence immunotherapy sensitivity. Nevertheless, the enrichment of low LIMD1 expression in responders suggests that LIMD1 loss contributes meaningfully to immune checkpoint regulation and highlights the need to consider LIMD1 status as part of a broader predictive framework for ICIs.

### Clinical and Biological Implications

Our study advances current understanding of immune checkpoint regulation by revealing a post-translational mechanism involving LIMD1 and ARIH1. The clinical relevance of this mechanism lies in its potential to stratify patients for immunotherapy and to guide the development of combination therapies. For example, targeting PD-L1 stabilisation pathways in LIMD1-deficient tumours may enhance the efficacy of ICIs, particularly in patients with intrinsic resistance. Furthermore, our previous work demonstrating proof-of-concept strategies for targeting LIMD1-deficient cells^53^ raises the exciting possibility of combining LIMD1-targeted therapies with ICIs to improve outcomes in NSCLC.

### Future Directions

While our findings provide compelling mechanistic and functional insights, further studies are needed to validate these observations *in vivo* and to explore the broader implications of LIMD1 loss across tumour types. Additionally, investigating LIMD1’s interaction with other immune checkpoints, such as PD-L2, or its role in modulating the tumour immune microenvironment will be critical for understanding its full therapeutic potential.

## Conclusion

In summary, our study establishes LIMD1 as a novel regulator of PD-L1 stability through its interaction with ARIH1, identifying a post-translational mechanism of immune evasion in NSCLC. By demonstrating that LIMD1 loss drives PD-L1 stabilisation and influences ICI response, we provide new insights into the molecular determinants of immune checkpoint regulation. These findings have significant implications for patient stratification and the development of targeted combination therapies, offering a new avenue for improving immunotherapy outcomes in LIMD1-deficient tumours.

## Acknowledgements and Contributions

We extend our gratitude to the members of our research teams for their invaluable contributions to this study. Tyson V. Sharp (TVS), the corresponding author, conceived and led the study, provided the foundational ideas, strategic direction, and supervision, and secured all associated funding. TVS also oversaw the data analysis and ensured the project’s alignment with its objectives.

The in vitro and molecular experiments were conducted by Kunal M. Shah (KMS), Paul T. Kennedy (PTK), Maria F. Contreras-Gerenas (MFCG), Kirsten Brooksbank (KB) and Oliver Yuan (OY), with strategic oversight and supervision provided by TVS. Data curation and analysis of the TRACERx and CPTAC datasets were performed by James R. M. Black (JRM) and Paul Grevitt (PG), under the guidance of Nicholas McGranahan (NM), Dimitris Lagos (DL), and TVS. For CPI response data, Krupa Thakkar (KT) acquired patient response data from various cohorts and performed gene expression analysis under the supervision of Kevin Litchfield (KL).

The manuscript was drafted by TVS, KMS, and PTK, with all authors contributing to subsequent editorial revisions. We acknowledge the technical assistance provided by the BCI Flow Cytometry Core Facility.

## Materials and Methods

### Cell culture

A549 cells were cultured in Dulbecco’s modified Eagle’s Medium (DMEM) (Gibco) supplemented with 10% v/v fetal bovine serum (Gibco) and penicillin/streptomycin. H1299, H358M, H3122, H2591, H28, CT26 and RCC48 cells were cultured in RPMI 1640. Medium (Gibco) supplemented with 10% v/v fetal bovine serum (Gibco) and penicillin/streptomycin. SAEC cells were cultured in small airway epithelial cell growth medium (PromoCell) with the designated supplements. Cells were cultured in humidified incubators at 37°C and 5% CO2. Recombinant Interferon-gamma was purchased from Millipore (IF002).

### Lentiviral transduction

A549, H1299 and RCC48 lentiviral lines were produced according to methods previously published ^38,54^.

### CRISPR-Cas9 gene editing

A549, H1299 LIMD1 knockout cells were generated with the Edit-R Cas9 nuclease (Horizon Discovery) and crRNAs targeting LIMD1 exon 1. CT26 *Limd1^-/-^*cells were generated by transfection with gRNAs targeting *Limd1* exon 1 following stable expression of Cas9-GFP. pLenti-Cas9-GFP was a gift from David Sabatini (Addgene plasmid # 86145).

### siRNA transfection

siRNAs were purchased from Horizon Discovery (On-TargetPLUS SmartPOOL), and Merck (Mission predesigned siRNA) and the siRNA panel for PD-L1 regulators was purchased from QIAGEN. Cells were transfected using DharmaFECT 1 transfection reagent (Horizon Discovery) or Lipofectamine RNAiMax (Life Technologies) according to the manufacturer’s instructions.

### PD-L1 3’UTR Luciferase reporter assay

psiCheck2 reporter for PD-L1 3’UTR is described elsewhere ^15^. Cells were transfected with siRNAs at 30 nM final concentration and 48h later transfected with psiCheck2 empty vector (VO) or PD-L1 3’ UTR. 24h post-transfection, cells were lysed in 1X Passive Lysis buffer (Promega) and 10 μl of lysate was transferred to white 96 well microplates for Dual Luciferase Assay (Promega), according to the manufacturer’s instructions. Luciferase activities were measured on the BMG FLUOstar Omega plate reader.

### Western blotting

Samples were prepared using RIPA buffer (150 mM NaCl, 50 mM Tris-HCl pH 8, 1 mM EDTA, 0.1% SDS, 1% NP-40, 0.5% sodium deoxycholate) containing protease and phosphatase inhibitors. Protein concentrations were determined with BCA and lysates were diluted in 5X SDS sample buffer with β-mercaptoethanol. Information on antibodies used is given in supplementary table 1. After SDS-PAGE, proteins were transferred onto PVDF membranes (Merck) and blocked using 5% non-fat dried milk in PBS-Tween 20. Following overnight incubation in the primary antibody, membranes were washed and incubated in HRP-conjugated secondary antibody before detection using Immobilon ECL substrate (Millipore) and imaged on Amersham Imagequant 600 (GE Healthcare). Densitometry was performed with ImageJ software on both the protein of interest bands and corresponding bands for loading control proteins.

### Immunoprecipitation

Cells grown in 15 cm dishes were lysed on ice in NP-40 lysis buffer (150 mM NaCl, 50 mM Tris pH 8, 0.7% v/v NP-40, 5% v/v glycerol) with protease and phosphatase inhibitors (Pierce, Roche). Lysates were cleared by centrifugation at 14,000 x g for 5 min at 4°C and then supernatants were transferred to new tubes. Input samples were prepared and then lysates were incubated with antibody overnight at 4°C. The following day, Protein G Mag Sepharose beads (Cytiva) were equilibrated in NP-40 buffer by two washes and then added to lysates. Beads were allowed to incubate with samples at room temperature for 1-2 h with rotation and then four washes with NP-40 buffer were performed. Proteins were eluted from beads by incubation for 5 min with 0.1M glycine pH 2.5. Elutions were transferred to new tubes, neutralised with 1M Tris-HCl pH 8 and diluted and boiled with 5X SDS sample buffer. Input and elution samples were subject to SDS-PAGE and transferred to PVDF membranes for immunoblotting.

For immunoprecipitation of HA-PD-L1 (pCMV-HA-PD-L1, a gift from Mien-Chie Hung, Addgene plasmid #121492) from A549 CRISPR-edited cells, plasmid DNA transfection was performed with Attractene Reagent (QIAGEN). 24h post-transfection, cells were lysed in NP-40 lysis buffer with protease and phosphatase inhibitors, lysates were cleared by centrifugation at 14,000 x g for 5 min at 4°C and then supernatants were transferred to new tubes. Input samples were prepared and cleared lysates were incubated with anti-HA magnetic beads (Pierce) overnight with gentle agitation at 4°C. The following day, beads were washed five times in RIPA buffer and proteins were eluted from the beads by boiling in 2.5X SDS sample buffer. Input samples and eluates were subsequently subjected to SDS-PAGE and immunoblotting.

### PD-L1 ubiquitination assay

Cells grown in 10 cm dishes were treated with 2 ng/mL IFN-γ overnight and then for 6 h with 10 μM MG132. Cells were lysed in RIPA buffer containing 10 mM N-ethylmaleimide. Lysates were cleared by centrifugation at 14,000 x g for 5 min at 4°C and then supernatants were transferred to new tubes. Input samples were prepared and then lysates were incubated overnight at 4°C with anti-PD-L1 antibody (ab213524, Abcam) or rabbit IgG isotype control (Cell Signalling Technologies). The following day, protein A Dynabeads (Life Technologies) were equilibrated in NP-40 lysis buffer and added to lysates to capture antibody-antigen complexes for 1h at room temperature with gentle agitation. Beads were then washed with RIPA buffer once, four times with high salt RIPA buffer (600 mM NaCl), and once more with RIPA buffer. Proteins were eluted from beads by boiling with 1X SDS sample buffer. Input and elution samples were subjected to SDS-PAGE and transferred to PVDF membranes for immunoblotting for ubiquitin and PD-L1.

### Flow cytometry

For analysis of surface PD-L1 expression in adherent cancer cell lines, wells were washed once with PBS before detaching cells with Accutase^®^ (STEMCELL Technologies). Single-cell suspensions were washed in PBS-1%FCS before staining with APC-conjugated anti-human-PD-L1 (29E.2A3, BioLegend), anti-mouse-PD-L1 (10F.9G2, BioLegend) or respective IgG controls. For analysis of T cell CD25 expression, suspension cells were collected, washed and stained with APC-conjugated anti-CD8 (SK1, BioLegend), PerCP-conjugated anti-CD4 (OKT4, BioLegend) and BV605™-conjugated anti-CD25 (BC96, BioLegend). Cells were stained for 30 min on ice and washed in PBS before analysis using FACSAria II (BD). FCS files were analysed using FlowJo V10.7 (TreeStar).

### Real-time quantitative PCR

Total RNA was extracted from cells using TRI reagent (Sigma) according to the manufacturer’s instructions. After determining concentration with a Nanodrop spectrophotometer, 1 µg of RNA was treated with amplification grade DNase I (Sigma) and diluted in nuclease-free water. Serial dilutions for RNA standard curves were prepared and these along with samples and no template controls, were used with GoTaq® 1-Step RT-qPCR System (Promega) to assay PD-L1 and beta-actin in duplicate reactions. PD-L1 Forward primer: 5’-TGG CAT TTG CTG AAC GCA TTT-3’, PD-L1 Reverse primer: 5’-AGT GCA GCC AGG TCT AAT TGT-3’. Beta-actin Forward primer: 5’-CTG GAA CGG TGA AGG TGA CA-3’, Beta-actin Reverse primer: 5’-AAG GGA CTT CCT GTA ACA ATG CA-3’. qPCR was performed on Applied Biosystems QuantStudio 5 real-time PCR machine and data was analysed using Design and Analysis software (Applied BioSystems).

### PBMC co-culture

96 well plates were coated with Ultra-LEAF™ anti-CD3 (OKT3, BioLegend) diluted in PBS. Plates were sealed and incubated at 37°C for 2 hours. PBMCs were harvested, centrifuged and resuspended in RPMI-1640-10% FCS growth media supplemented with recombinant human interleukin-2 (10 ng/mL, Roche) before incubation at 37°C 5% CO_2_ for 3 days. 3×10^3^ target A549 cells were seeded into each well and incubated for 5 hours or until adhered. Anti-CD3-activated PBMCs were harvested from 96 well plates and resuspended to 15×10^4^ cells/mL in complete media. A549-H2B-mCherry target cells expressing a nuclear fluorescent reporter protein were seeded in growth media containing hIL-2 (final concentration 10 ng/mL). For studies of immune checkpoint function, cultures were treated with pembrolizumab (anti-PD-1, Merck), atezolizumab (anti-PD-L1, Genentech) or humanised IgG1 Control (InVivoMAb, BioXcell) at 10µg/mL. 50×10^3^ PBMCs were then seeded into cultures at predetermined target to effector ratios (E: T), to a final 200 µL culture volume. Images were captured using the IncuCyte® ZOOM live cell imaging system (Essen Biosciences).

### 3D culture/spheroid experiments

For 3D spheroid assays, 500 viable target A549-mKate cells expressing a cytosolic reporter protein were seeded into ultra-low attachment (ULA) plates (Corning) and cultured for 7 days until spheroids had formed. Cell culture media was removed carefully and 1×10^5^ activated PBMCs were seeded into wells containing spheroids. Images were captured using the IncuCyte® S3 live cell imaging system (Essen Biosciences). For measurement of cell death within co-cultures, media was supplemented with CellTox Green™ reagent (Promega), with red and green fluorescence measured at 650 nm and 530 nm, respectively.

### TRACERx Data Analysis

TRACERx gene expression and clinico-pathological data was obtained from Martinez-Ruiz et al 2023 ^55^. LIMD1 copy number information was obtained from Frankell et al., 2023 ^34^., Tumours were considered as having ‘Clonal’ loss if there was evidence of copy number loss relative to ploidy throughout the LIMD1 locus in all constituent regions of a tumour. Tumours were considered as having ‘Subclonal’ loss if there was evidence of copy number loss relative to ploidy anywhere in the LIMD1 locus in at least one constituent region of a tumour, as well as evidence of a non-loss copy number state relative to ploidy anywhere in the LIMD1 locus in at least one constituent region of a tumour. Tumours were considered as having ‘No loss’ if there was no evidence of copy number loss relative to ploidy at any point of the LIMD1 locus in any of the constituent regions of a tumour. Gene expression data was VST-transformed for downstream analysis of the impact of LIMD1 gene expression. CD8 T-cell estimates were obtained using the approach described in Danaher et al., 2017^56^; and were accessed from Puttick et al., 2024^57^. Scores estimating the expression of the Buffa et al., ^39^ genes were obtained by calculating the median VST-transformed gene expression value for constituent genes. Where indicated, Wilcoxon tests performed in this work are two-sided, using the function wilcox.test() in base R. For correlation analyses for Buffa hypoxia score and CD8+ T cell score, the P value was calculated using a linear mixed-effects model with tumour as the random variable.

### CPI Data Analysis

Three cohorts were used for the immune checkpoint blockade analysis – 26 patients from the Samsung Medical Centre ^58^, 323 patients from the OAK clinical trial ^43^ and 57 from the POPLAR trial ^44^. All patients are treated with anti-PD1 or anti-PDL1 monotherapy. Clinical data such as treatment type was collected, and response outcomes were harmonised using the RECIST criteria ^59^.

RNA-sequencing data was derived from the studies in a FASTQ format and then processed through the RIMA pipeline ^60^. Gene quantification was conducted using tximport (Salmon) and was TPM normalised for downstream analysis.

---

**Supplementary Figure 1.** (**A**) VST-transformed LIMD1 expression inversely correlates with an hypoxic gene expression signature in LUAD and LUSC. (**B**) CD274 expression positively correlates with an hypoxic gene signature in LUAD but not in LUSC.

**Supplementary Figure 2.** (**A**) Immunoblots of PD-L1 in lung adenocarcinoma cells transfected with LIMD1-targeting or scrambled control siRNA (20nM) 48 hours before stimulating with IFNγ (2 ng/mL) overnight. Vinculin was probed as a loading control (**B**) Stable LIMD1 over-expression in MCA-205 fibrosarcoma cells. (**C**) Flow cytometric analysis of surface PD-L1 in MCA-205. (**D**) Ajuba knockdown in mesothelioma cell lines upregulates PD-L1 protein. H2591 and H28 cell lines were transfected with Scr negative control siRNA or Ajuba siRNA and treated with or without IFNγ. Whole-cell lysates were immunoblotted for PD-L1, Ajuba and vinculin as a loading control.

**Supplementary Figure 3.** A549 Cas9 control and LIMD1^-/-^ clone 5 were treated with increasing concentrations of recombinant IFNγ overnight and then harvested for whole cell lysate for immunoblotting **(A)** or total RNA for RT-qPCR of PD-L1 **(B)**. Graphs in A show densitometry for PD-L1 intensity normalised to loading control. (**B**) The graph shows mean PD-L1 mRNA levels normalised to beta-actin mRNA levels for five independent experiments. Error bars denote SEM. No statistically significant difference in PD-L1 mRNA was seen between Cas9 and LIMD1^-/-^ clone 5 according to two-way ANOVA with Sidak’s multiple comparisons test.

**Supplementary Figure 4.** (**A**) PD-L1 3’UTR reporter assay. A549 cells were transfected with the indicated siRNAs at 30 nM and after 48h were transfected with psiCHECK2 empty vector (VO) or psiCHECK2-PD-L1 3’UTR. Cells were harvested 24h later and lysates assayed for firefly and Renilla luciferase activities. Data shown mean normalised Renilla luciferase activity for PD-L1 3’UTR relative to VO for at least three independent experiments +/- SEM. * denotes p<0.05 according to Mann Whitney u-test; ns is not significant. (**B**) A549 Cas9 and LIMD1^-/-^ cells (clone 5 and clone 10) were treated with vehicle or chloroquine (CQ) at 40 µM overnight. Whole-cell lysates were immunoblotted for PD-L1, LIMD1, LC3B and Vinculin as a loading control. Graph shows mean fold change in PD-L1 levels relative to vehicle treatment for each line for three independent experiments. Ns means not significant according to the student’s t-test. (**C**) Representative western blots for PD-L1 and vinculin for siRNA transfections in A549 Cas9 and LIMD1^-/-^ C5 and C10 clones for main figure 4D.

**Supplementary Figure 5.** Microscopy-based measurement of cytotoxicity within atezolizumab-treated A549-PBMC 3D spheroid co-cultures. (**A**) Representative end-point images show nuclei density at 175 hours (scale bar = 250 μm). (**B**) Data are mean ± sem for a representative PBMC donor. PBMCs were pre-activated with anti-CD3 (2 µg/mL for 3 days), and cultures were treated with IgG control, pembrolizumab or atezolizumab (10 µg/mL for 7 days).

