## Supplementary figures and images for "LIMD1 Loss as an Early Driver of PD-L1 Upregulation and Immune Evasion in Lung Cancer"

### Supp Fig 1

##### Supplementary Figure 1

A

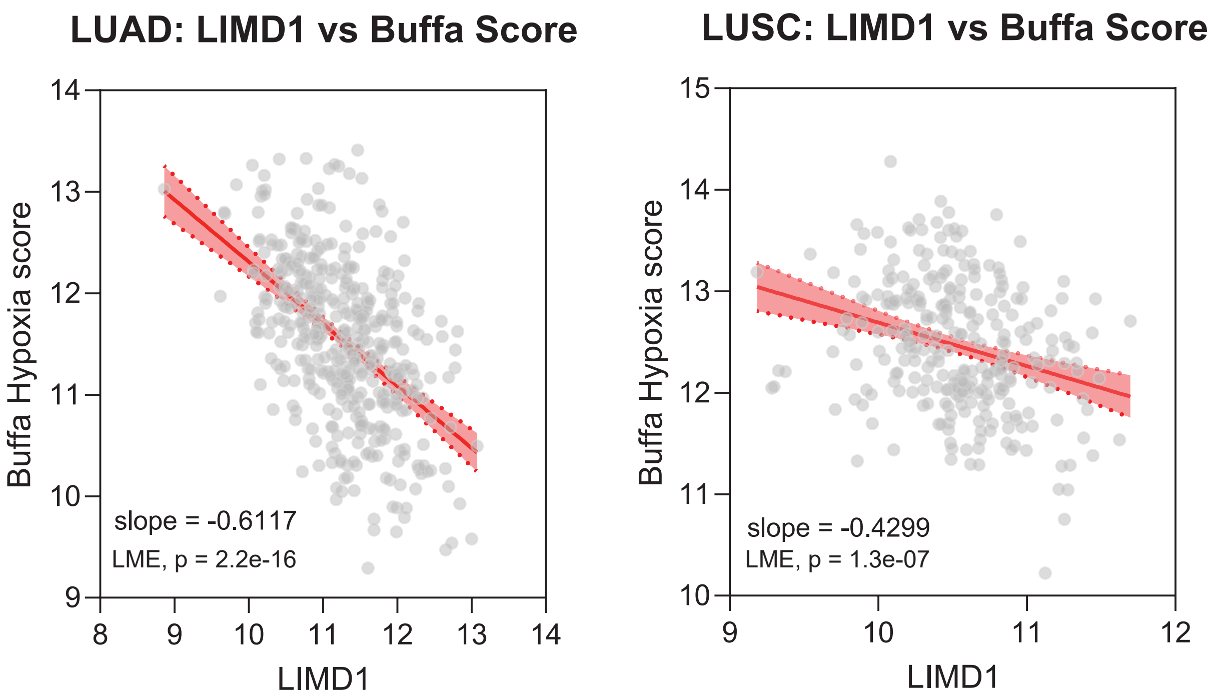


B

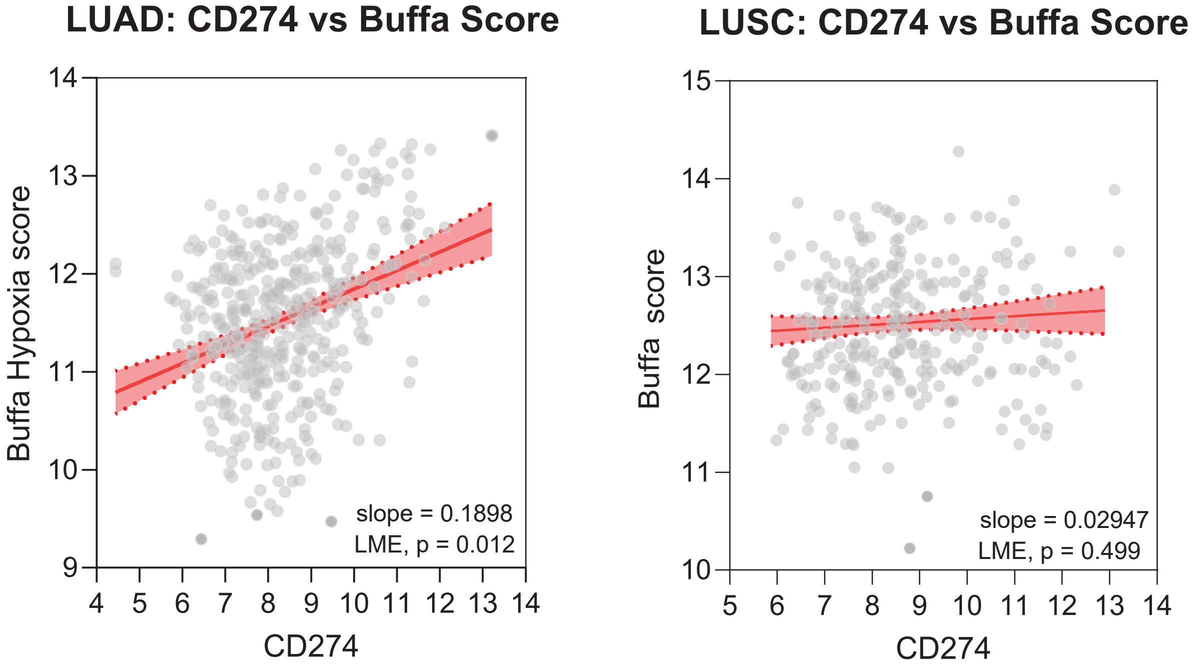

### Supp Fig 2

#### Supplementary Figure 2

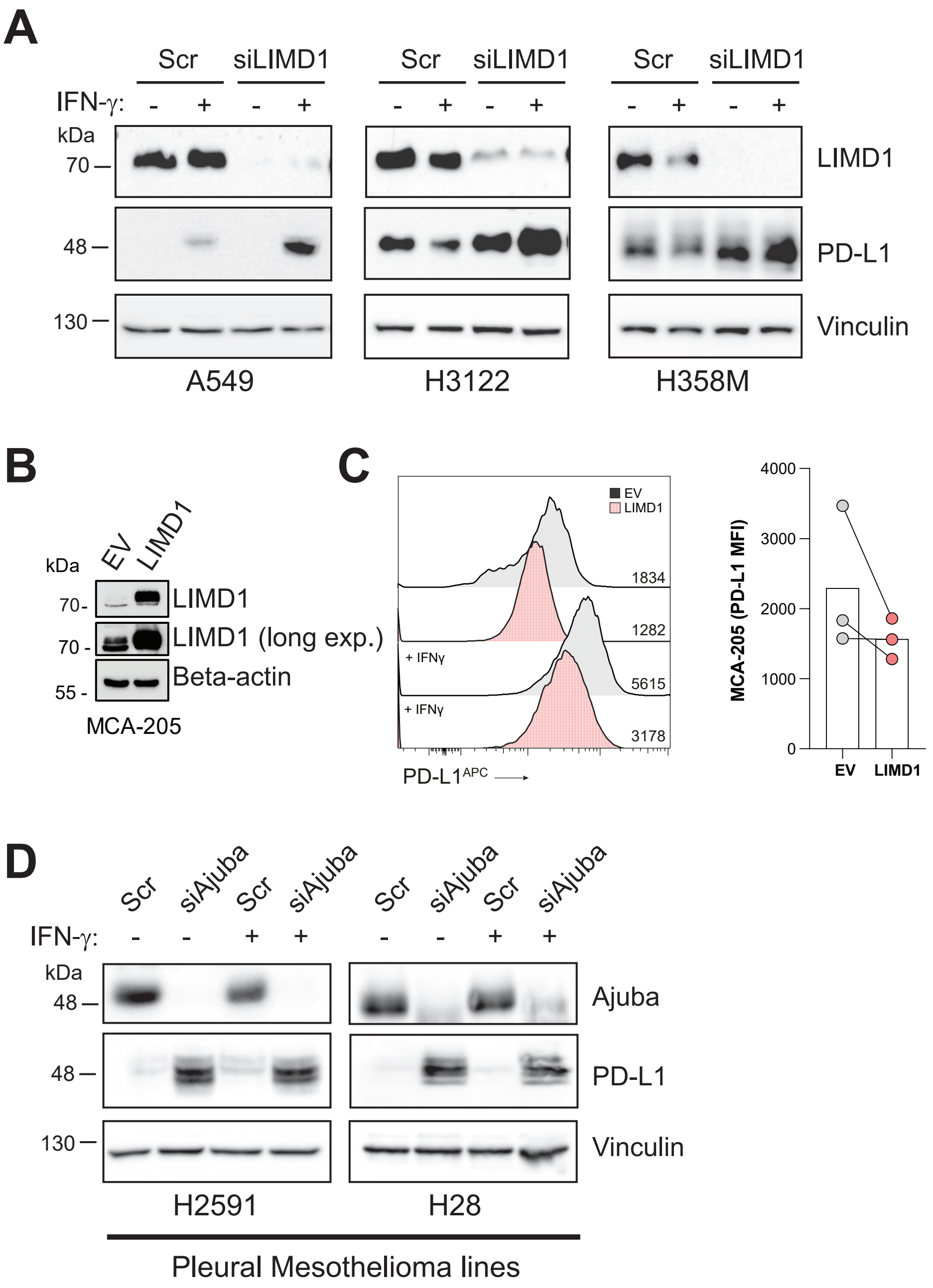

### Supp Fig 3

#### Supplementary Figure 3

A

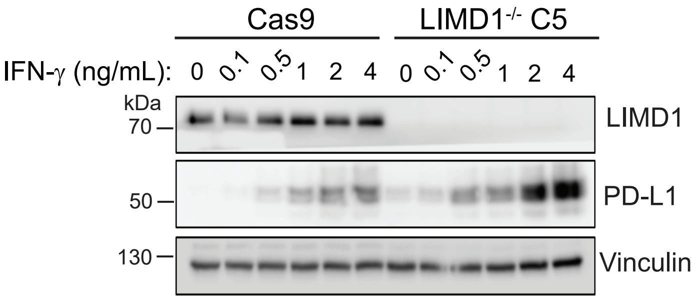


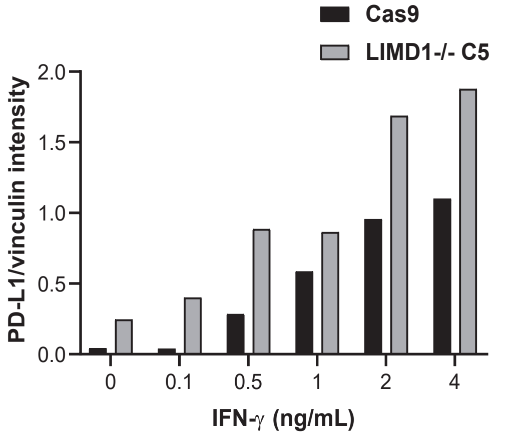


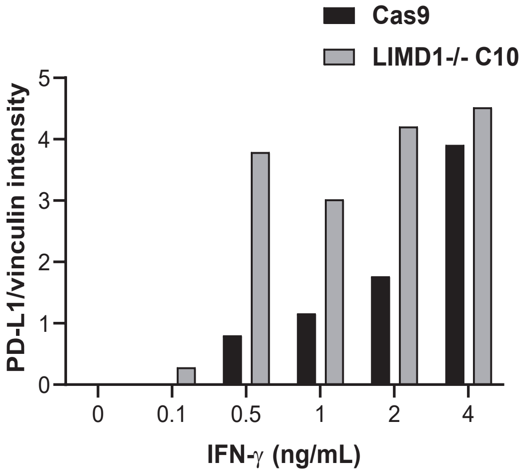


B

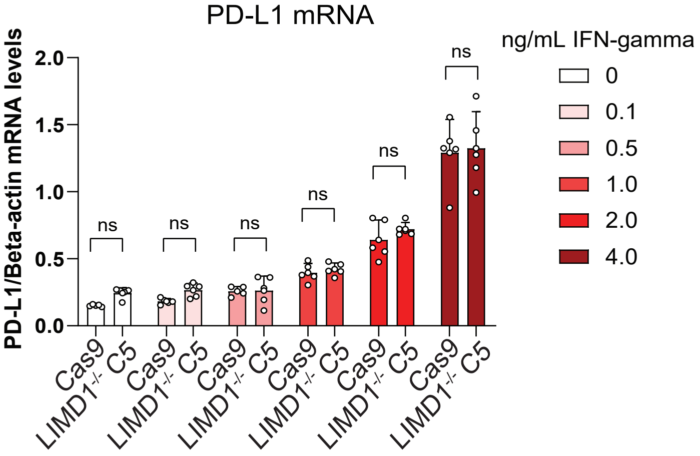

### Supp Fig 4

##### Supplementary Figure 4

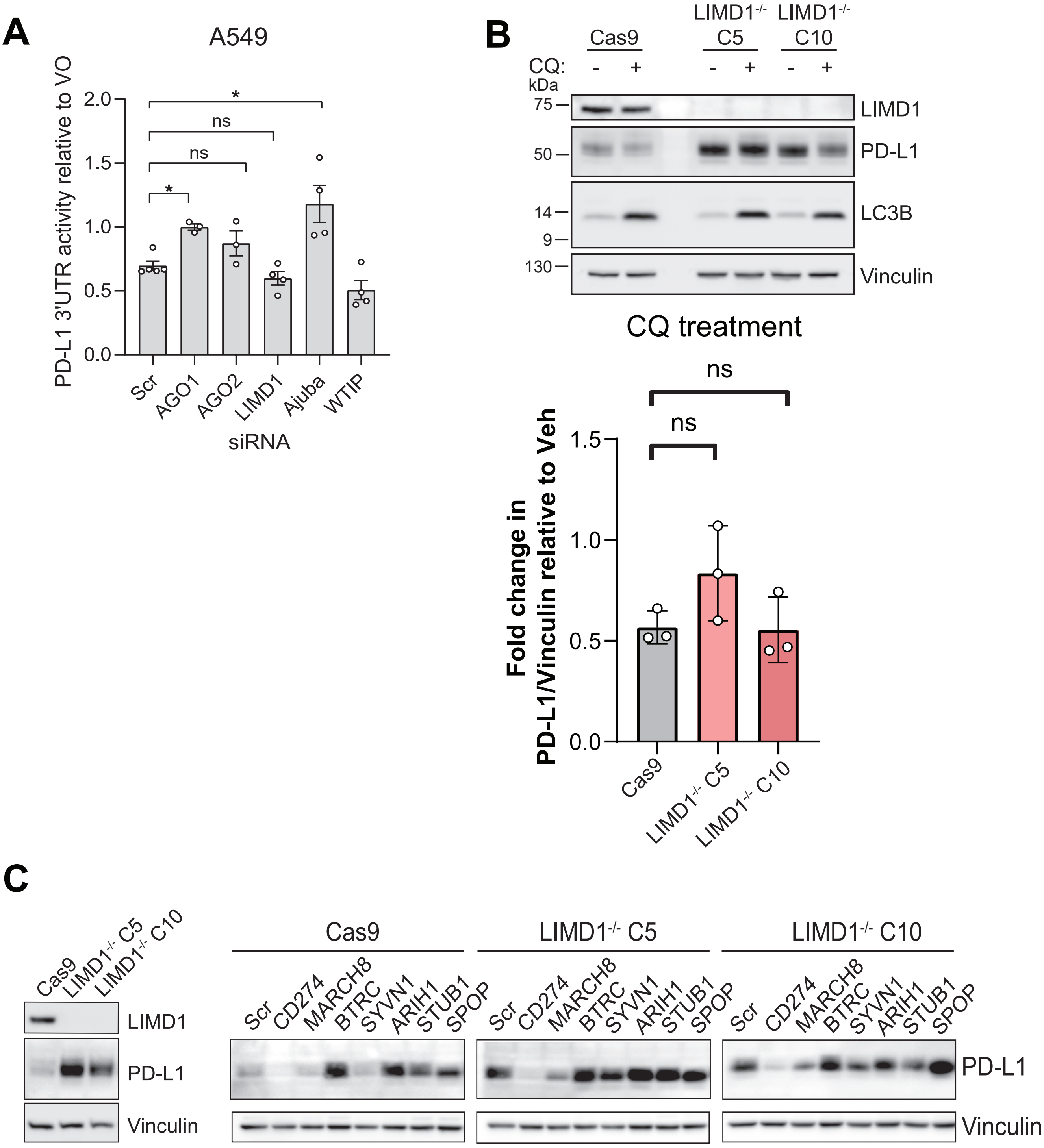

### Supp Fig 5

##### Supplementary Figure 5

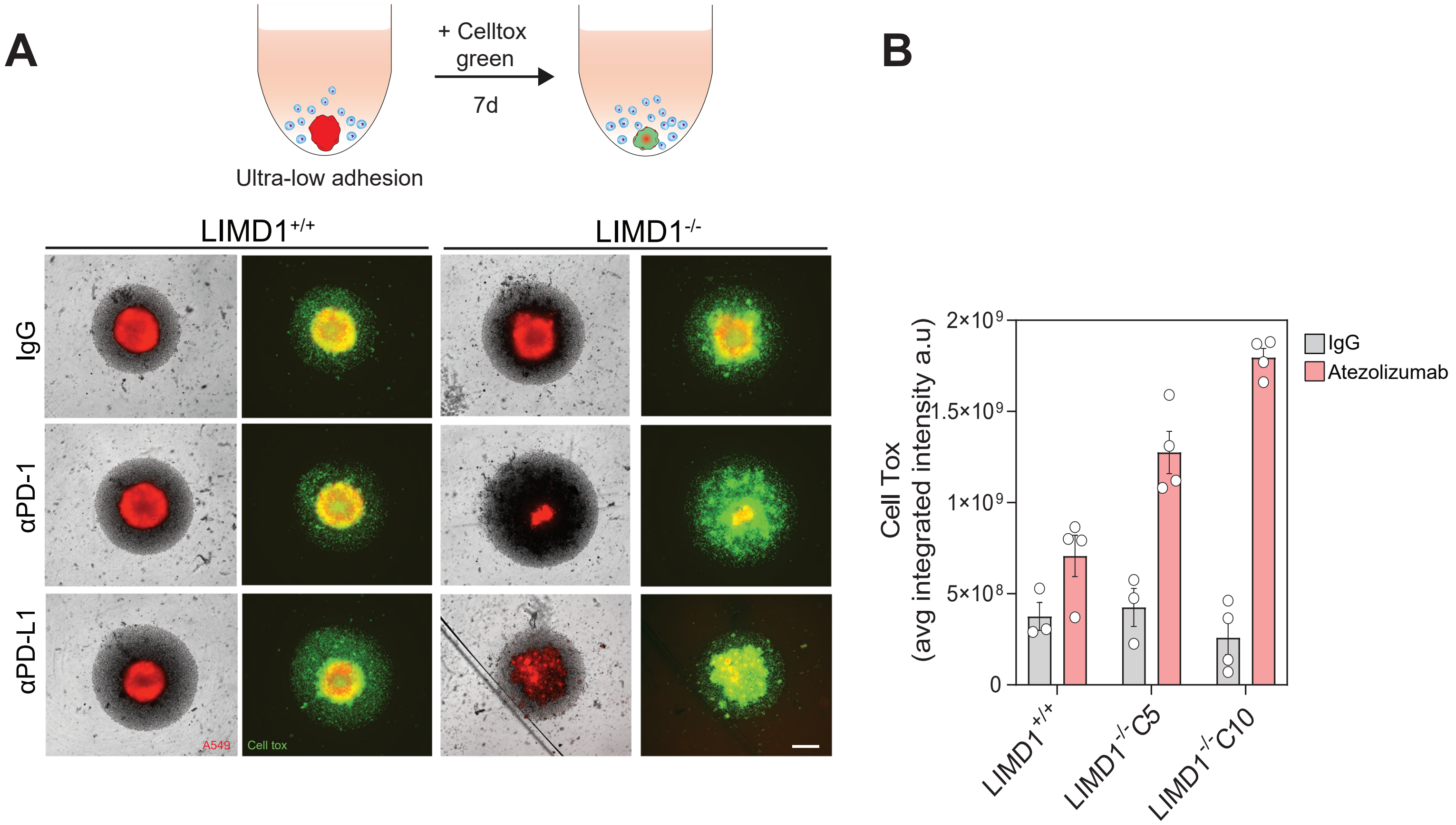
